# Ancestral Origin of the CXCL17–GPR25 System Traced to the Lobe-Finned Fish *Latimeria chalumnae*

**DOI:** 10.1101/2025.09.16.676457

**Authors:** Jie Yu, Juan-Juan Wang, Wen-Feng Hu, Ya-Li Liu, Zeng-Guang Xu, Zhan-Yun Guo

## Abstract

The roles of C-X-C motif chemokine ligand 17 (CXCL17) and its receptor, G protein-coupled receptor 25 (GPR25), in immune regulation are gaining recognition, but their evolutionary origins and phylogenetic distributions have yet to be determined. In this study, we identified a CXCL17 gene in the lobe-finned fish Latimeria chalumnae (coelacanth), a close living relative of tetrapods. This gene produces two alternatively spliced secretory proteins, designated Lc-CXCL17a and Lc-CXCL17b, which display limited sequence similarity to known CXCL17s from mammals or ray-finned fishes. Recombinant forms of Lc-CXCL17a and Lc-CXCL17b were generated via bacterial overexpression, purification, and enzymatic processing. Functional assays, including NanoLuc Binary Technology (NanoBiT)-based β-arrestin recruitment, ligand–receptor binding, and chemotaxis assays, demonstrated that both proteins tightly bound to and potently activated coelacanth GPR25 in a manner dependent on their C-terminal residues, because removal of these residues markedly diminished activities. These findings provide the first evidence that a functional CXCL17–GPR25 system exists in lobe-finned fishes, supporting the hypothesis that this ligand–receptor pair originated in ancient fishes and was transmitted to vertebrate lineages through lobe-finned fish ancestors. This work broadens understanding of chemokine evolution and highlights the coelacanth as a valuable model for tracing conserved immune signaling pathways.

## 1. Introduction

C-X-C motif chemokine ligand 17 (CXCL17) is a chemoattractant predominantly expressed in mucosal tissues such as the lung, stomach, salivary gland, and esophagus, as reported in the Human Protein Atlas (https://www.proteinatlas.org). It recruits various immune cells, including T cells, monocytes, macrophages, and dendritic cells, to these tissues [1-14]. Beyond its role in mucosal immunity, CXCL17 has also been implicated in tumor development, likely through the regulation of tumor-associated immune responses [15-21]. The chemoattractant function of CXCL17 is presumed to be mediated by a plasma membrane receptor; however, the identity of this receptor has long been debated. A decade ago, CXCL17 was initially described as an agonist of the orphan G protein-coupled receptor 35 (GPR35) [22], but this finding could not be reproduced by other groups [23-25]. More recently, CXCL17 was proposed to act as a modulator of the chemokine receptor CXCR4 [25], although this claim also awaits independent confirmation. In addition, CXCL17 has been reported to activate the MAS-related receptor MRGPRX2 [26]. Our recent work demonstrated that human CXCL17 indeed activates MRGPRX2 at micromolar concentrations, while also activating two other MAS-related receptors, MRGPRX1 and MAS1, with slightly lower efficiency [27]. Notably, activation of these MAS-related receptors occurs independently of CXCL17’s conserved C-terminal fragment [27], indicating that they are unlikely to represent its evolutionarily conserved receptor. Most recently, Ocón’s group and our team independently showed that CXCL17 can activate the poorly characterized orphan G protein-coupled receptor 25 (GPR25) via its conserved C-terminal fragment [28,29], strongly suggesting that GPR25 is the evolutionarily conserved receptor for CXCL17.

GPR25 is an A-class G protein-coupled receptor (GPCR) that is predominantly expressed in immune cells, including T cells, plasma cells, and B cells, according to the Human Protein Atlas (https://www.proteinatlas.org). GPR25 orthologs are broadly conserved from fishes to mammals, indicating that this receptor fulfills important functions across vertebrates. In contrast, the existence of CXCL17 orthologs in non-mammalian vertebrates remained unclear until our recent identification of CXCL17 and a CXCL17-like variant in zebrafish (*Danio rerio*) and several other ray-finned fishes [30]. These fish CXCL17 orthologs represent previously uncharacterized proteins with unknown functions, as they share little overall amino acid sequence similarity with mammalian CXCL17s and therefore cannot be reliably detected using conventional sequence-based searches [30].

In our recent study [31], we found that human CXCL17 can efficiently activate the GPR25 ortholog from the lobe-finned fish *Latimeria chalumnae* (coelacanth), suggesting that a CXCL17 ortholog might also be present in this iconic “living fossil” fish. As one of the closest extant fish relatives of tetrapods, *L. chalumnae* has long been of particular interest in evolutionary research. In this study, we identified a candidate *CXCL17* gene in *L. chalumnae* based on its genomic location and the characteristics of its encoded proteins. Through alternative splicing, this gene generates two small secretory proteins differing by a single amino acid, which we designated as Lc-CXCL17a and Lc-CXCL17b. These proteins show no significant overall amino acid sequence similarity to known mammalian or fish CXCL17s, and no additional homologs were retrieved from public databases, suggesting that they are unique and may preserve ancestral features. Recombinant Lc-CXCL17a and Lc-CXCL17b exhibited strong activity toward coelacanth GPR25 (Lc-GPR25) in NanoLuc Binary Technology (NanoBiT)-based β-arrestin recruitment, ligand–receptor binding, and chemotaxis assays, but this activity was almost abolished upon deletion of four C-terminal residues. These findings indicate that Lc-CXCL17a and Lc-CXCL17b function as endogenous agonists of Lc-GPR25, supporting the hypothesis that the CXCL17–GPR25 pair originated in ancient fishes and was passed on to vertebrate lineages through lobe-finned fish ancestors. This work provides new insights into the evolutionary origin and distribution of the CXCL17–GPR25 system in vertebrates.

## 2. Materials and methods

### 2.1. Preparation of the recombinant coelacanth CXCL17s

The DNA fragment encoding the N-terminally 6×His-tagged mature peptide of Lc-CXCL17a was chemically synthesized and ligated into a pET vector via Gibson assembly, resulting in the expression construct pET/6×His-Lc-CXCL17a. The expression construct for 6×His-Lc-CXCL17b was generated via the QuikChange approach using pET/6×His-Lc-CXCL17a as the mutagenesis template. To generate the expression constructs encoding the C-terminally 3×Arg-tagged version, the coding region of 6×His-Lc-CXCL17a and 6×His-Lc-CXCL17b were amplified by polymerase chain reaction (PCR) and ligated into the pET vector via Gibson assembly, resulting in the expression constructs pET/6×His-Lc-CXCL17a-3×Arg and pET/6×His-Lc-CXCL17b-3×Arg (Fig. S1). The coding region of the N-terminally SmBiT-fused version was PCR amplified using pET/6×His-Lc-CXCL17a-3×Arg as the template and ligated into the pET vector via Gibson assembly, resulting in the expression construct pET/6×His-SmBiT-Lc-CXCL17a-3×Arg (Fig. S1). The coding region of the coelacanth CXCL17s in these expression constructs was confirmed by DNA sequencing.

Recombinant expression of the coelacanth CXCL17s in *Escherichia coli* was conducted according to our previous procedure developed for human or zebrafish CXCL17s [29,30]. After the transformed BL21(DE3) cells were induced by isopropyl-β-D-thiogalactopyranoside (IPTG), expression of the coelacanth CXCL17s was detected by sodium dodecyl sulfate-polyacrylamide gel electrophoresis (SDS-PAGE). If the expected protein was successfully overexpressed, the bacteria were lysed by sonication and the inclusion bodies were solubilized via an *S*-sulfonation approach. Thereafter, the coelacanth CXCL17 precursors were purified via a Ni^2+^ column, subjected to *in vitro* refolding, and further purified by high performance liquid chromatography (HPLC) using a C_18_ reverse-phase column (Zorbax 300SB-C18, 9.4 × 250 mm, Agilent Technologies, Santa Clara, CA, USA). The eluted fractions were manually collected, lyophilized, and treated with recombinant carboxypeptidase B (Yaxin Bio, Shanghai, China) to remove three C-terminal Arg residues. After further purification by HPLC, the recombinant coelacanth CXCL17s with mature C-terminus were dissolved in 1.0 mM aqueous hydrochloride (pH 3.0) and quantified by ultra-violet absorbance at 280 nm according to their extinction coefficient (16500 M^-1^ cm^-1^ for the 6×His-Lc-CXCL17a, 6×His-Lc-CXCL17b, and the C-terminally truncated mutant; 17990 M^-1^ cm^-1^ for 6×His-SmBiT-Lc-CXCL17a).

### 2.2. Generation of the expression constructs for the coelacanth GPR25

The expression constructs for coelacanth GPR25 (Lc-GPR25) were generated in our recent study [31]. The doxycycline (Dox)-inducible expression constructs PB-TRE/Lc-GPR25 and PB-TRE/sLgBiT-Lc-GPR25 encode an untagged Lc-GPR25 or an N-terminally secretory large NanoLuc fragment (sLgBiT)-fused Lc-GPR25, respectively. The Dox-inducible expression construct pTRE3G-BI/Lc-GPR25-LgBiT:SmBiT-ARRB2 coexpresses a C-terminally LgBiT-fused Lc-GPR25 and an N-terminally SmBiT-fused human β-arrestin 2 (SmBiT-ARRB2) via a bidirectional promoter.

### 2.3. The NanoBiT-based β-arrestin recruitment assays

The NanoBiT-based β-arrestin recruitment assays were conducted according to our previous procedure [29-31]. Briefly, the coexpression construct pTRE3G-BI/Lc-GPR25-LgBiT:SmBiT-ARRB2 and the expression control vector pCMV-Tet3G (Clontech, Mountain View, CA, USA) were cotransfected into human embryonic kidney (HEK) 293T cells using the transfection reagent Lipo8000 (Beyotime Technology, Shanghai, China). Next day, the cells were trypsinized, seeded into white opaque 96-well plates, and cultured in induction medium (complete DMEM plus 1.0 ng/mL of Dox) for ∼24 h to ∼90% confluence. To conduct the β-arrestin recruitment assay, the induction medium was removed and pre-warmed activation solution (serum-free DMEM plus 1% bovine serum albumin) was added (40 μL/well, containing 0.5 μL of NanoLuc substrate stock from Promega, Madison, WI, USA), and bioluminescence data were immediately collected for ∼4 min on a SpectraMax iD3 plate reader (Molecular Devices, Sunnyvale, CA, USA). Subsequently, recombinant coelacanth CXCL17s (diluted in the activation solution) was added (10 μL/well), and bioluminescence data were continuously collected for ∼10 min. The measured absolute bioluminescence signals were corrected for inter well variability by forcing all curves after addition of NanoLuc substrate (without ligand) to same level and plotted using the SigmaPlot 10.0 software (SYSTAT software, Chicago, IL, USA). To obtain the dose-response curve, the measured bioluminescence data at highest point were plotted with the agonist concentrations using the SigmaPlot 10.0 software (SYSTAT software).

### 2.4. The NanoBiT-based ligand-receptor binding assays

The NanoBiT-based ligand-receptor binding assays were developed using sLgBiT-Lc-GPR25 as a receptor source and 6×His-SmBiT-Lc-CXCL17a as a tracer according to our previous procedures developed for some other GPCRs [30, 32-34]. Briefly, HEK293T cells were transiently transfected with the expression construct PB-TRE/sLgBiT-Lc-GPR25 with or without cotransfection of the human tyrosylprotein sulfotransferases coexpression construct pTRE3G-BI/TPST1:TPST2. Next day, the transfected cells were trypsinized, seeded into white opaque 96-well plates, and cultured in induction medium (complete DMEM plus 20 ng/mL of Dox) for ∼24 h to ∼90% confluence. To conduct the binding assays, the induction medium was removed and pre-warmed binding solution (serum-free DMEM plus 0.1% bovine serum albumin and 0.01% Tween-20) was added (50 μL/well). For saturation binding assays, the binding solution contains different concentrations of 6×His-SmBiT-Lc-CXCL17a. For competition binding assays, the binding solution contains a constant concentration of 6×His-SmBiT-Lc-CXCL17a and different concentrations of competitors.

To measure cell surface expression level of sLgBiT-Lc-GPR25, binding solution contains 80 nM of synthetic HiBiT peptide. After incubation at room temperature for ∼1 h, diluted NanoLuc substrate (30-fold dilution in the binding solution) was added (10 μL/well), and bioluminescence was immediately measured on a SpectraMax iD3 plate reader (Molecular Devices). The measured bioluminescence data were expressed as mean ± standard deviation (SD, *n* = 3) and fitted to one-site binging model via the SigmaPlot 10.0 software (SYSTAT software).

### 2.5. Chemotaxis assays

The transwell chemotaxis assays were conducted according to our previous procedure [29-31]. Briefly, HEK293T cells were transfected with the expression construct PB-TRE/Lc-GPR25 using the transfection reagent Lipo8000 (Beyotime Technology). Next day, the transfected cells were changed to the induction medium (complete DMEM plus 1.0 ng/mL of Dox) and continuously cultured for ∼24 h. Thereafter, the cells were trypsinized, suspended in serum-free DMEM at the density of ∼5×10^5^ cells/mL, and seeded into polyethylene terephthalate membrane (8 μm pore size)-coated permeable transwell inserts that were pretreated with serum-free DMEM. The inserts were then put into a 24-well plate containing chemotactic agent (WT or truncated coelacanth CXCL17 diluted in serum-free DMEM plus 0.2% bovine serum albumin, 500 μL/well). After cultured at 37°C for ∼5 h, solution in the inserts were removed and cells on the upper face of the permeable membrane were wiped off using cotton swaps, and cells on the lower face of the permeable membrane were fixed with 4% paraformaldehyde solution, stained with crystal violet staining solution (Beyotime Technology), and observed under an Olympus APX100 microscope (Tokyo, Japan). The migrated cells were quantitated using the ImageJ software and the results were expressed as mean ± SD (*n* = 3).

## 3. Results

### 3.1. Identification of possible CXCL17 orthologs from the lobe-finned fish L. chalumnae

To search for potential CXCL17 orthologs in the lobe-finned fish *L. chalumnae*, we retrieved all of its secretory proteins with fewer than 200 amino acids from the UniProt database (https://www.uniprot.org/uniprotkb?query=Latimeria+chalumnae&facets=proteins_with%3A49%2Clength%3A%5B1+TO+200%5D). Among 320 candidate proteins, none exhibited the key sequence features of CXCL17 orthologs, namely a C-terminal Xaa-Pro-Yaa motif (with Xaa and Yaa typically being large aliphatic residues) and six conserved cysteine residues in the mature peptide.

In humans, the *CXCL17* gene (gene ID: 284340) is located near *CEACAM8*, *CEACAM1*, *LIPE*, *CNFN*, *MEGF8*, *TMEM145*, *PRR19*, *PAFAH1B3*, and *CIC* (Fig. S2). Based on this syntenic context, we hypothesized that the coelacanth *CXCL17* gene would be located adjacent to one of these markers. Consistent with this prediction, analysis of the published *L. chalumnae* genome in the NCBI database identified a candidate gene (gene ID: 106704074) on chromosome 25 near *MEGF8* (Fig. S3). Alternative splicing of this locus produces two uncharacterized small secretory proteins, which we designated as Lc-CXCL17a and Lc-CXCL17b (Fig. 1A and Table S1).

**Fig. 1.**
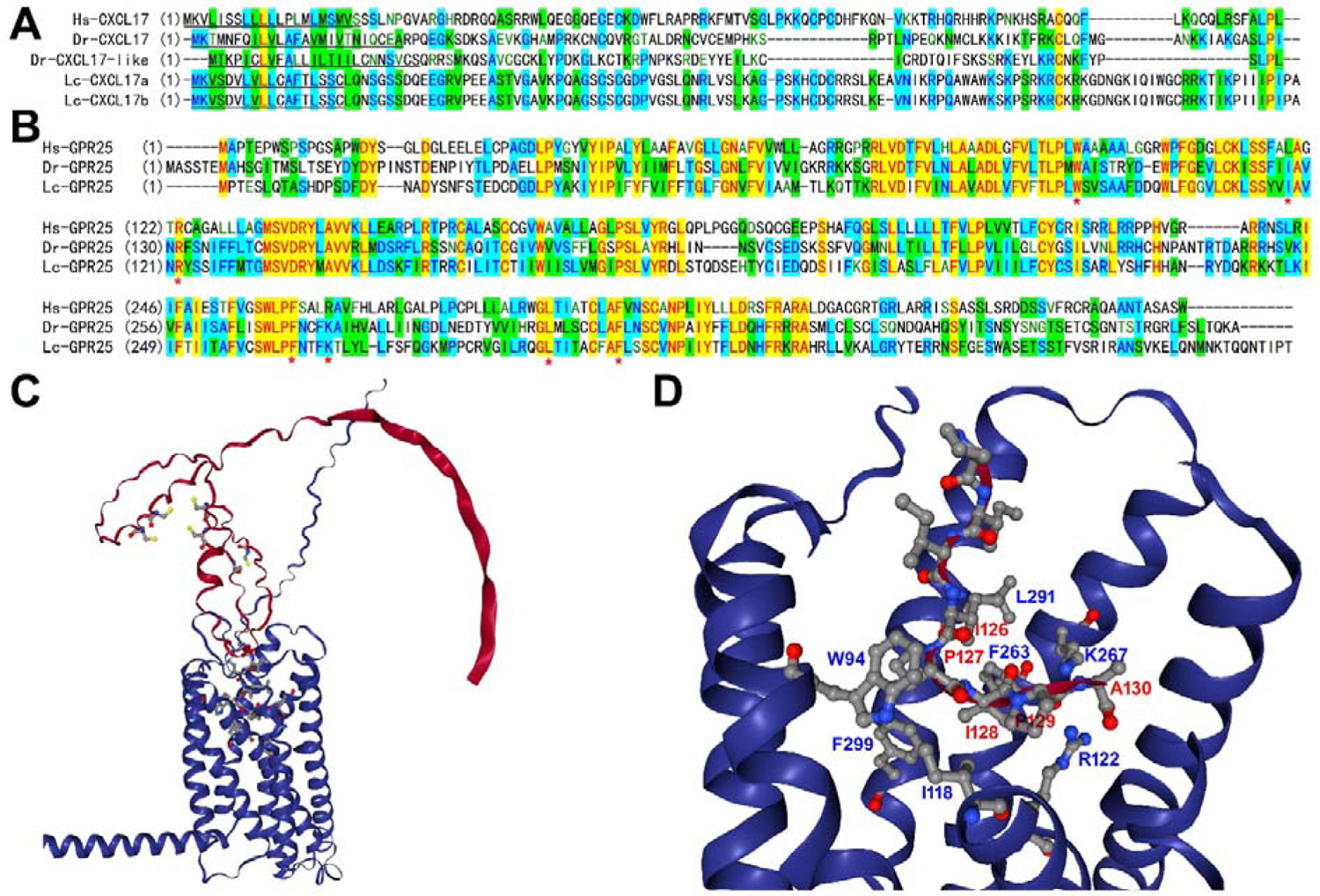
Identification of possible CXCL17 orthologs from the lobe-finned fish *L. chalumnae*. (**A,B**) Amino acid sequence alignment of the CXCL17 orthologs (**A**) or GPR25 orthologs (**B**) from human, zebrafish, and coelacanth. Their information is downloaded from the NCBI gene database and listed in Table S1. The predicted signal peptide of CXCL17 orthologs is underlined. These sequences were aligned via AlignX algorithm using the Vector NTI 11.5.1 software. In panel B, the receptor residues involving ligand binding are indicated by red asterisks. (**C,D**), Overall view (C) and close-up view (D) of the AlphaFold3-predicted structure of mature Lc-CXCL17a binding to Lc-GPR25. The binding structure was predicted via the online AlphaFold3 server (https://alphafoldserver.com) and viewed by an online server (https://nglviewer.org/ngl). In the predicted structure, Lc-CXCL17a is shown in red, and Lc-GPR25 in blue. The six Cys residues and eight C-terminal residues of Lc-CXCL17a are shown as sticks-and-balls. The receptor residues interacting with Lc-CXCL17a are shown as sticks-and-balls and labeled. For clarity, some fragments of Lc-GPR25 are not shown in panel D. The predicted possible interactions are summarized in Table S2.

Both Lc-CXCL17a and Lc-CXCL17b contain a predicted N-terminal signal peptide, and their mature forms differ by only a single amino acid (Fig. 1A; Table S1). Each contains six cysteine residues arranged in a CXC-CXC-C-C pattern, identical to that found in mammalian CXCL17s. However, sequence alignment revealed that coelacanth CXCL17s share no significant overall similarity with human CXCL17 (Hs-CXCL17) or the newly identified zebrafish orthologs (Dr-CXCL17 and Dr-CXCL17-like), explaining why they had not been recognized previously (Fig. 1A). In contrast, coelacanth GPR25 (Lc-GPR25) shows strong sequence homology with its human and zebrafish counterparts (Fig. 1B), indicating that GPR25 orthologs are much more conserved during evolution than CXCL17 orthologs. The mature Lc-CXCL17 proteins have predicted isoelectric points of ∼10.6, suggesting they are positively charged under physiological conditions.

This property is consistent with their putative chemoattractant role, as positively charged chemokines can interact with negatively charged glycosaminoglycans on the cell surface to establish concentration gradients that guide immune cell migration.

No homologous proteins were retrieved from public databases using BLAST searches with Lc-CXCL17a or Lc-CXCL17b (https://blast.ncbi.nlm.nih.gov), supporting the idea that these proteins are unique and may retain ancient features. Notably, both proteins contain a distinctive C-terminal hydrophobic stretch (PIIIPIPA) composed primarily of proline and isoleucine residues, which includes two partially overlapping Xaa-Pro-Yaa motifs (Fig. 1A).

Structural predictions generated with the AlphaFold3 algorithm suggest that mature Lc-CXCL17a and Lc-CXCL17b adopt highly flexible conformations with low overall prediction confidence. Nonetheless, AlphaFold3 predicted its binding to Lc-GPR25 with ipTM values of ∼0.6. In these binding models, the hydrophobic C-terminal segment of Lc-CXCL17a inserts into the orthosteric ligand-binding pocket of Lc-GPR25 (Fig. 1C), with the final five residues forming extensive interactions with conserved receptor residues (Fig. 1D and Table S2). For instance, the negatively charged C-terminal carboxyl group of Lc-CXCL17a forms an electrostatic interaction with the guanidinium group of the receptor’s R122, while the last but three proline of Lc-CXCL17a engages in hydrophobic interactions with W94 of the receptor (Fig. 1D; Table S2). Because Lc-CXCL17b differs by only a single amino acid in the central region of the peptide, it is predicted to display nearly identical interaction patterns with Lc-GPR25 (Table S2). The residues involved in ligand binding are conserved across GPR25 orthologs (Fig. 1B), lending strong support to the reliability of these predicted binding models.

### 3.2. Preparation of the coelacanth CXCL17s via bacterial overexpression

To rapidly obtain coelacanth CXCL17s for functional assays, we employed bacterial overexpression. However, the N-terminally 6×His-tagged mature forms of Lc-CXCL17a and Lc-CXCL17b were not expressed in *Escherichia coli*, likely due to their toxicity to host cells. To overcome this limitation, we introduced three additional Arg residues at their C-terminus. These residues could later be conveniently removed by carboxypeptidase B following purification and refolding. As expected, the modified constructs (6×His-Lc-CXCL17a-3×Arg and 6×His-Lc-CXCL17b-3×Arg) were successfully overexpressed in *E. coli*, forming inclusion bodies as confirmed by SDS-PAGE (Fig. 2A, B). After solubilization of inclusion bodies using an *S*-sulfonation method, the 6×His-tagged precursors were purified via Ni²-affinity chromatography. SDS-PAGE analysis of the eluted fractions revealed the expected monomeric precursor along with higher-molecular-weight oligomers (Fig. 2A, B), indicating that the coelacanth CXCL17s tend to undergo intermolecular cross-linking. HPLC analysis demonstrated that both recombinant proteins exhibited a broad major peak, observed both before and after *in vitro* refolding (Fig. 2C, D). SDS-PAGE analysis of the refolded samples (Fig. 2C, D, inner panels) confirmed the presence of the expected monomer (marked with an asterisk), together with larger oligomers and smaller degradation products, no matter presence or absence of the reducing reagent dithiothreitol (DTT). Subsequent enzymatic treatment with carboxypeptidase B efficiently removed the three C-terminal Arg residues. Following this maturation step, HPLC analysis revealed a broad peak with a longer retention time (Fig. 2C, D, green trace), and SDS-PAGE analysis confirmed that the monomer was the predominant species within this fraction (Fig. 2C, D, inner panels). Together, these results demonstrated that coelacanth CXCL17s can be effectively produced through bacterial overexpression of C-terminally Arg-tagged precursors, followed by enzymatic maturation with carboxypeptidase B.

**Fig. 2.**
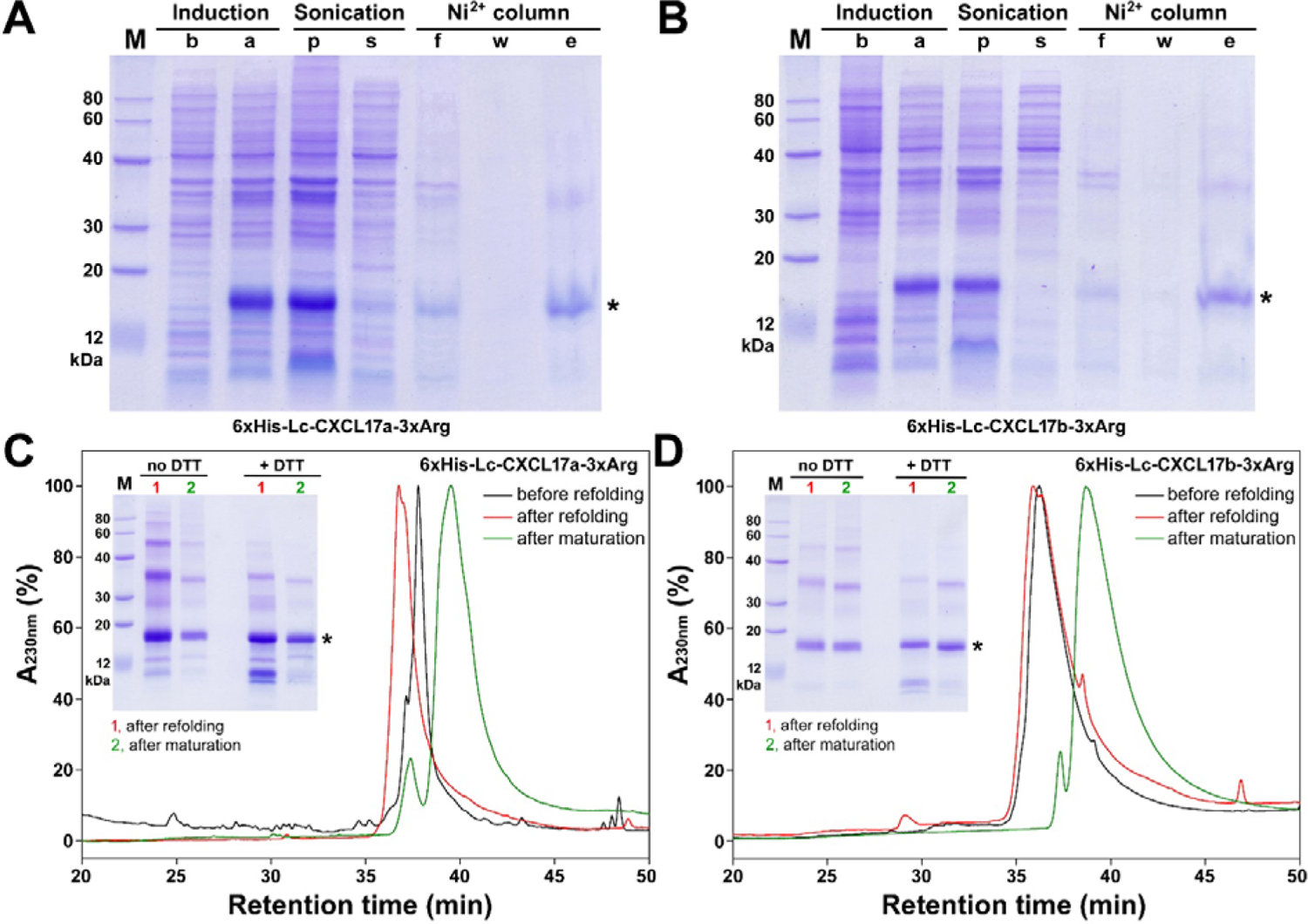
Preparation of the recombinant coelacanth CXCL17s. (**A,B**) SDS-PAGE analysis of the samples of 6×His-Lc-CXCL17a-3×Arg (**A**) or 6×His-Lc-CXCL17b-3×Arg (**B**) at different stages. Lane (M), protein ladder; lane (b), before IPTG induction; lane (a), after IPTG induction; lane (p), pellet after sonification; lane (s), supernatant after sonification; lane (f), flowthrough from the Ni^2+^ column; lane (w), washing fraction by 30 mM imidazole; lane (e), eluted fraction by 250 mM imidazole. After electrophoresis, the SDS-gel was stained by Coomassie brilliant blue R250. Band of the monomeric Lc-CXCL17 precursor was indicated by an asterisk. (**C,D**) HPLC analysis of the samples of 6×His-Lc-CXCL17-3×Arg (**C**) or 6×His-Lc-CXCL17-3×Arg (**D**) at different stages. **Inner panel**, SDS-PAGE analysis of Lc-CXCL17 samples at different stages. Lane (M), protein ladder; lane (1), after refolding; lane (2), after maturation by carboxypeptidase B. The SDS-gel was stained by Coomassie brilliant blue R250 after electrophoresis. Band of the monomeric Lc-CXCL17 was indicated by an asterisk.

### 3.3. Activation of the coelacanth GPR25 by the recombinant coelacanth CXCL17s

To determine whether coelacanth CXCL17s are capable of activating coelacanth GPR25, we performed NanoBiT-based β-arrestin recruitment assays. In this system, a C-terminally large NanoLuc fragment (LgBiT)-fused Lc-GPR25 (Lc-GPR25-LgBiT) was coexpressed with an N-terminally low-affinity complementation tag (SmBiT)-fused human β-arrestin 2 (SmBiT-ARRB2) in HEK293T cells. Upon agonist stimulation, activated Lc-GPR25-LgBiT recruits SmBiT-ARRB2, resulting in a proximity-driven reconstitution of NanoLuc activity and a corresponding bioluminescence signal. Because endogenous receptors lack the LgBiT fusion, this assay is unlikely to be influenced by endogenous receptors.

Following induction of Lc-GPR25-LgBiT and SmBiT-ARRB2 expression, the addition of NanoLuc substrate yielded only low baseline bioluminescence (Fig. 3A, B). Subsequent treatment with recombinant 6×His-Lc-CXCL17a or 6×His-Lc-CXCL17b rapidly increased bioluminescence in a dose-dependent manner (Fig. 3A, B), demonstrating that both proteins effectively activate Lc-GPR25 and promote β-arrestin recruitment. Dose–response analysis (Fig. 3C) yielded EC-values of approximately 70 nM for 6×His-Lc-CXCL17a and 40 nM for 6×His-Lc-CXCL17b. By contrast, the C-terminally truncated mutant, 6×His-Lc-[desC4]CXCL17a, exhibited markedly reduced activity, with concentrations up to 1.0 μM producing only minimal bioluminescence changes (Fig. 3D). These findings indicate that coelacanth CXCL17 orthologs efficiently activate Lc-GPR25 through their C-terminal residues, suggesting that CXCL17 employs a conserved mechanism to activate GPR25 that dates back to ancient fishes.

**Fig. 3.**
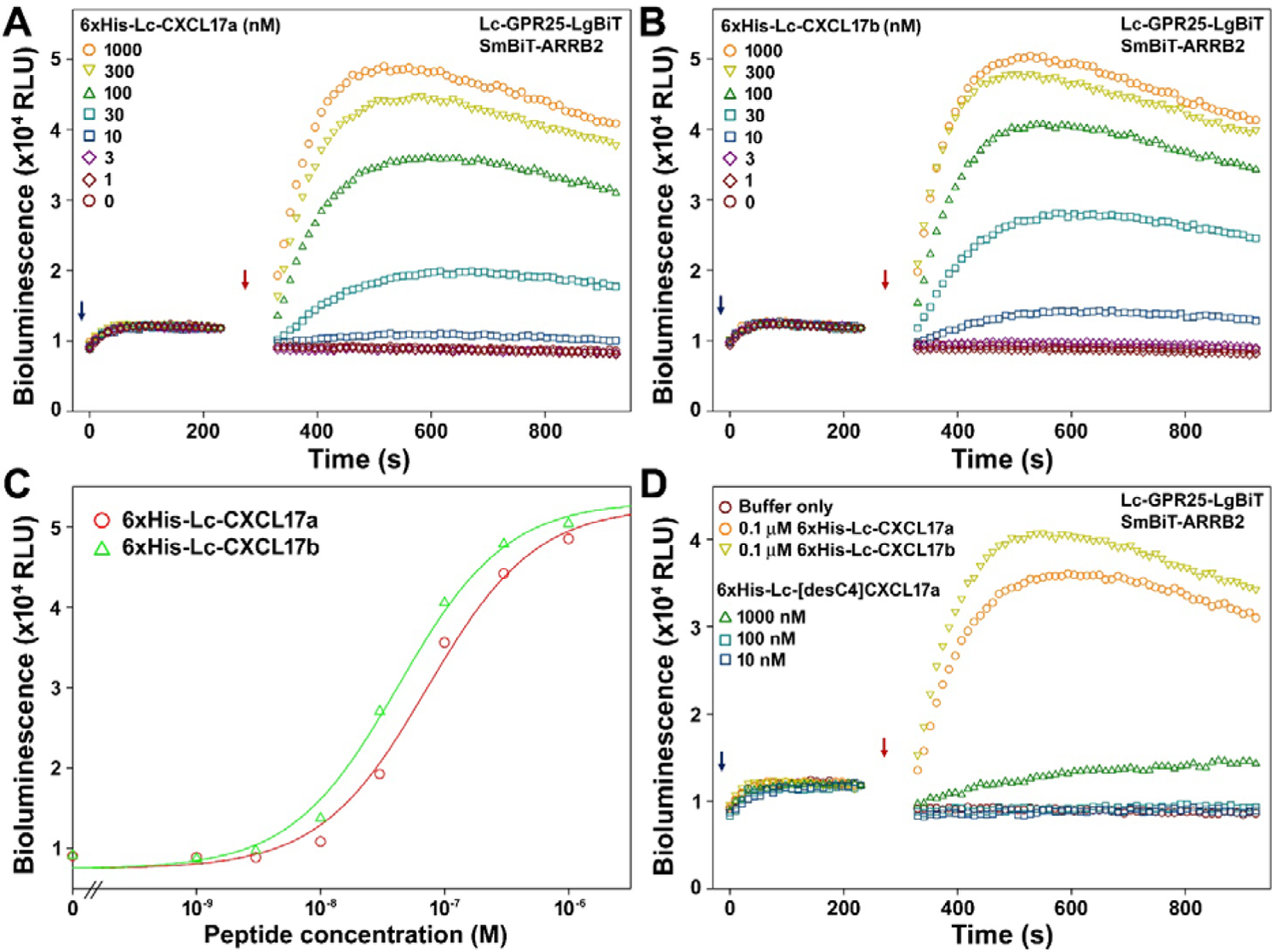
NanoBiT-based β-arrestin recruitment assays of the coelacanth GPR25 induced by the recombinant coelacanth CXCL17s. (**A,B**) Effect of the WT 6×His-Lc-CXCL17a (**A**) or 6×His-Lc-CXCL17b (**B**). (**C**) Dose curve of the WT coelacanth CXCL17s activating Lc-GPR25 assayed as β-arrestin recruitment in panel A and B. (**D**) Effect of the C-terminally truncated 6×His-Lc-[desC4]CXCL17a. In these assays, NanoLuc substrate and different concentrations of WT or mutant coelacanth CXCL17 were sequentially added to living HEK293T cells coexpressing Lc-GPR25-LgBiT and SmBiT-ARRB2, and bioluminescence data were continuously measured on a plate reader. Typical bioluminescence change profiles are shown in panel A, B, and D, and the calculated dose response curves are shown in panel C. The blue arrows indicate addition of NanoLuc substrate, and red arrows indicate addition of the coelacanth CXCL17.

### 3.4. Binding of the recombinant coelacanth CXCL17s with the coelacanth GPR25

To evaluate direct binding between coelacanth CXCL17s and coelacanth GPR25, we employed a NanoBiT-based homogeneous binding assay. In this system, a secretory LgBiT (sLgBiT) was fused to the extracellular N-terminus of Lc-GPR25, and a SmBiT tag was fused to the N-terminus of Lc-CXCL17a. Binding of recombinant 6×His-SmBiT-Lc-CXCL17a to sLgBiT-Lc-GPR25 would bring the two fragments into proximity, leading to luciferase reconstitution. Owing to the requirement for engineered fusion tags, this assay is highly specific and unlikely to be affected by endogenous receptors.

Following overexpression, purification, refolding, and carboxypeptidase B treatment, the binding tracer 6×His-SmBiT-Lc-CXCL17a was obtained. Its activity was confirmed via the NanoBiT-based β-arrestin recruitment assay, where it induced a rapid, dose-dependent increase in bioluminescence (Fig. 4A), with an EC- of ∼100 nM (Fig. 4A, inner panel), comparable to that of 6×His-Lc-CXCL17a and 6×His-Lc-CXCL17b.

**Fig. 4.**
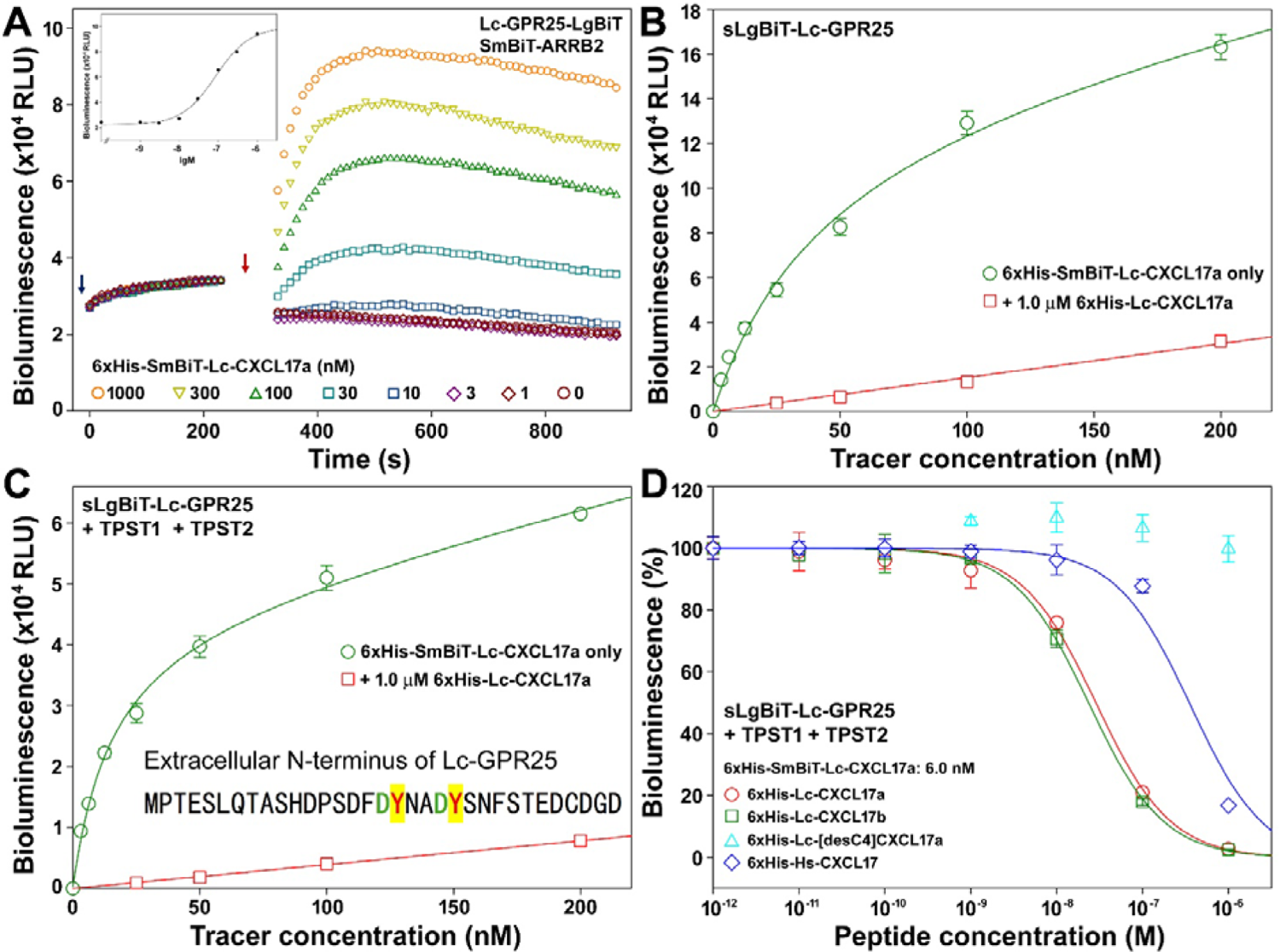
NanoBiT-based homogenous binding assays of the recombinant coelacanth CXCL17s with coelacanth GPR25. (**A**) Activity of the recombinant 6×His-SmBiT-Lc-CXCL17a towards Lc-GPR25 assayed as NanoBiT-based β-arrestin recruitment. NanoLuc substrate and different concentrations of 6×His-SmBiT-Lc-CXCL17 were sequentially added to living HEK293T cells coexpressing Lc-GPR25-LgBiT and SmBiT-ARRB2, and bioluminescence was continuously measured on a plate reader. The blue arrow indicates the addition of NanoLuc substrate, and the red arrow indicates the addition of 6×His-SmBiT-Lc-CXCL17a. The calculated dose response curve is shown in the inner panel. (**B,C**) Saturation binding of 6×His-SmBiT-Lc-CXCL17a with sLgBiT-Lc-GPR25 under absence (B) or presence (C) of human TPST1 and TPST2. The measured bioluminescence data are expressed as mean ± SD (*n* = 3) and plotted using the SigmaPlot10.0 software. Total binding data (green circles) were fitted with the function of Y = B_max_X/(K_d_+X) + k_non_X, the non-specific binding data (red squares) were fitted with linear curves. (**D**) Competition binding of some recombinant CXCL17s with Lc-GPR25 measured via the NanoBiT-based homogenous binding assays. The measured bioluminescence data are expressed as mean ± SD (*n* = 3) and fitted with sigmoidal curves using the SigmaPlot10.0 software.

Subsequently, we examined the binding of 6×His-SmBiT-Lc-CXCL17a to sLgBiT-Lc-GPR25 using saturation binding assays. When the tracer was added to living HEK293T cells expressing sLgBiT-Lc-GPR25, with or without coexpression of human tyrosylprotein sulfotransferases (TPST1 and TPST2), hyperbolic binding curves were obtained (Fig. 4B, C). The calculated dissociation constant (K_d_) was 45 ± 10 nM (*n* = 3) in the absence of TPST1 and TPST2, and 15.5 ± 1.7 nM (*n* = 3) in their presence, indicating that tyrosylprotein sulfotransferase coexpression substantially enhanced binding affinity. These results suggest that the coexpressed tyrosylprotein sulfotransferases likely catalyze sulfation of tyrosine residues at the extracellular N-terminus of Lc-GPR25 (Fig. 4C, inner panel), thereby facilitating stronger ligand–receptor interactions. Competition with 1.0 μM of 6×His-Lc-CXCL17a drastically reduced the bioluminescence signal (Fig. 4B, C), confirming the specificity of the tracer binding to Lc-GPR25.

Finally, we performed NanoBiT-based competition binding assays to compare the binding potencies of different CXCL17 proteins toward Lc-GPR25 (Fig. 4D). Both recombinant 6×His-Lc-CXCL17a and 6×His-Lc-CXCL17b displaced the tracer with similar efficiency, yielding IC-values of ∼30 nM. In contrast, recombinant human CXCL17 (6×His-Hs-CXCL17) displaced the tracer less effectively, with an IC- of ∼400 nM, indicating that its binding affinity for Lc-GPR25 was approximately 13-fold weaker than that of the coelacanth proteins. Notably, the truncated mutant 6×His-Lc-[desC4]CXCL17a failed to displace the tracer even at concentrations up to 1.0 μM, demonstrating a complete loss of binding to Lc-GPR25. This finding is consistent with its negligible activity in the β-arrestin recruitment assay.

### 3.5. Chemotactic effect of the recombinant coelacanth CXCL17s

The chemotactic activity of coelacanth CXCL17s was evaluated in transwell assays using HEK293T cells transfected to express Lc-GPR25. Recombinant 6×His-Lc-CXCL17a and 6×His-Lc-CXCL17b promoted cell migration in a dose-dependent manner, whereas the C-terminally truncated mutant 6×His-[desC4]CXCL17a exhibited no effect even at concentrations up to 1.0 μM (Fig. 5). These findings indicate that coelacanth CXCL17s act as chemoattractants by activating Lc-GPR25, supporting the conclusion that CXCL17 serves as an endogenous agonist of GPR25 in the lobe-finned fish *L. chalumnae*.

**Fig. 5.**
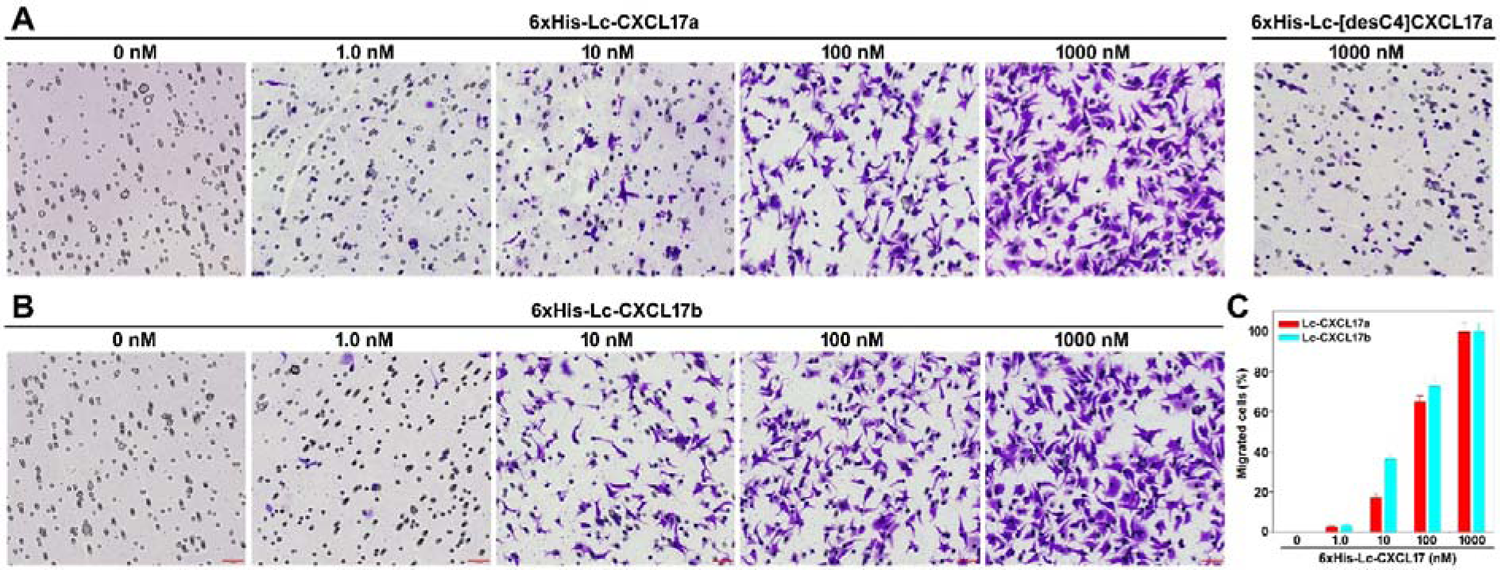
Chemotactic effect of the recombinant coelacanth CXCL17s. (**A**) Representative images of the migrated HEK293T cells expressing Lc-GPR25 after induced by WT or C-terminally truncated 6×His-CXCL17a. (**B**) Representative images of the migrated HEK293T cells expressing Lc-GPR25 after induced by WT 6×His-CXCL17b. (**C**) Quantitative analysis of the migrated HEK293T cells expressing Lc-GPR25 after induced by WT coelacanth CXCL17s. Transfected HEK293T cells expressing Lc-GPR25 were seeded into the permeable membrane-coated inserts and induced by chemotactic solution in the lower chamber. After the assay, cells on the upper face of the permeable membrane were wiped off, and cells on the lower face of the permeable membrane were fixed, stained, and observed under a microscope. Representative images of the migrated HEK293T cells are shown in panel A and B. The scale bar in these images is 50 μm. The migrated cells were quantitatively analyzed using the ImageJ software and the results are expressed as mean ± SD (*n* = 3) and shown in panel C.

## 4. Discussion

In this study, we identified CXCL17 orthologs in the lobe-finned fish *L. chalumnae*, one of the closest extant fish relatives of tetrapods. These findings suggest that CXCL17 originated in ancient fishes and was transmitted to other vertebrate lineages through lobe-finned fish ancestors. As a well-known “living fossil” fish, *L. chalumnae* closely resembles its prehistoric relatives. Fossil evidence indicates that coelacanths first appeared during the Devonian period, approximately 400 million years ago, flourished in the Triassic, and then declined markedly in the Late Cretaceous around 70 million years ago. Today, only two coelacanth species, *L. chalumnae* and *L. menadoensis*, persist in restricted regions of the Indian Ocean. Genomic analyses have demonstrated that the genes of *L. chalumnae* evolve at a significantly slower rate than those of tetrapods [35,36], suggesting that many of its proteins may retain ancestral characteristics. The coelacanth CXCL17s possess a distinctive hydrophobic C-terminal stretch (PIIIPIPA) containing two partially overlapping Xaa-Pro-Yaa motifs. Such features are absent from CXCL17s in extant mammals and ray-finned fishes but may have been widespread among ancient forms. Moreover, the organization of the *CXCL17* gene and its neighboring genes is conserved between humans and *L. chalumnae* (Fig. S2 and S3), supporting the notion of a shared evolutionary origin.

As demonstrated in our recent study, zebrafish and several other closely related ray-finned fishes possess two CXCL17 paralogs, CXCL17 and CXCL17-like [30]. In contrast, only a single *CXCL17* gene was identified in the lobe-finned fish *L. chalumnae*. As shown in Fig. S4, the zebrafish *cxcl17-like* gene is located near *ponzr3*, *ponzr4*, *ponzr5*, *ponzr6*, *ugt5g1*, and *cldn7a*; however, these neighboring genes appear to be absent in *L. chalumnae* according to the NCBI genome database. Alternatively, the zebrafish *cxcl17-like* gene is also positioned near *fgf11a*, *chrnb1*, and *tmem256* (Fig. S4). These flanking genes are present in the coelacanth genome (Fig. S5), yet no *cxcl17-like* gene was detected in this region. These findings suggest that the *cxcl17-like* gene likely arose in certain ray-finned fishes but may never have existed in lobe-finned lineages. Consequently, extant tetrapods, from amphibians to mammals, as descendants of ancient lobe-finned fishes, are also expected to lack the *cxcl17-li*ke gene in their genomes.

Our study identified a *CXCL17* gene in the extant lobe-finned fish *L. chalumnae*. It is therefore reasonable to infer that this gene was also present in ancient lobe-finned fishes and subsequently transmitted to tetrapods, including amphibians, reptiles, birds, and mammals. While CXCL17 orthologs have not yet been found in amphibians, reptiles, or birds, their existence is highly probable. Based on conserved genomic positioning (adjacent to marker genes such as *MEGF8*, *TMEM145*, *CNFN*, *CEACAM1*, or *LIPE*) and characteristic sequence features (a C-terminal Xaa-Pro-Yaa motif, mature peptides of fewer than 200 residues, and six conserved cysteine residues), it is likely that CXCL17 orthologs in these lineages will be discovered and functionally characterized in future studies.

## Data availability statement

The data of this study are available in this manuscript, as well as the associated supplementary information.

## CRediT Author Contribution

Conceptualization, Z.Y.G.; investigation, J.Y., J.J.W., and W.F.H.; writing – original draft, Z.Y.G.; writing – review & editing, Z.Y.G., J.Y., and J.J.W.; formal analysis, Y.L.L., and Z.G.X.; project administration, Y.L.L., and Z.G.X.; funding acquisition, Z.Y.G.; supervision, Z.Y.G. All authors read and approved the final manuscript.

## Abbreviations

ARRB2: human β-arrestin 2
CXCL17: C-X-C motif chemokine ligand 17
DMEM: Dulbecco’s modification of Eagle’s medium
Dox: doxycycline
DTT: dithiothreitol
GPCR: G protein-coupled receptor
GPR25: G protein-coupled receptor 25
HEK: human embryonic kidney
HiBiT: high-affinity complementation tag for NanoBiT
HPLC: high performance liquid chromatography
IPTG: isopropyl-β-D-thiogalactopyranoside
LgBiT: large NanoLuc fragment for NanoBiT
NanoBiT: NanoLuc Binary Technology
NanoLuc: nanoluciferase
NCBI: the National Center for Biotechnology Information
PCR: polymerase chain reaction
SD: standard deviation
SDS-PAGE: sodium dodecyl sulfate-polyacrylamide gel electrophoresis
sLgBiT: secretory LgBiT
SmBiT: low-affinity complementation tag for NanoBiT
WT: wild-type.

