## supplemental Table S1-S2 and Fig. S1-S5 for "Ancestral Origin of the CXCL17–GPR25 System Traced to the Lobe-Finned Fish *Latimeria chalumnae*"

**Contents:**

**Table S1.** Information about CXCL17 orthologs and GPR25 orthologs from human, zebrafish, or coelacanth.

**Table S2.** Summary of possible interactions of the coelacanth CXCL17a and CXCL17b with the coelacanth GPR25 predicted by AlphaFold3 algorithm.

**Fig. S1.** The nucleotide and amino acid sequence of the coelacanth CXCL17 precursors overexpressed in *E. coli*.

**Fig. S2.** Position of the human *CXCL17* gene in human genome.

**Fig. S3.** Position of the coelacanth *CXCL17* gene in the genome of *Latimeria chalumnae*.

**Fig. S4.** Position of the zebrafish *cxcl17-like* gene in the genome of *Danio rerio*.

**Fig. S5.** Genes near to *TMEM256* in the genome of *Latimeria chalumnae*.

**Table S1.** Information about CXCL17 orthologs and GPR25 orthologs from human, zebrafish, or coelacanth. The information is downloaded from the NCBI gene database (https://www.ncbi.nlm.nih.gov/gene). The predicted signal peptide of CXCL17 orthologs is shaded.

| Class | Name in this study | Species | Gene ID | mRNA ID | Protein ID | Amino acid sequence |
|---|---|---|---|---|---|---|
| CXCL17 orthologs | Hs-CXCL17 | *Homo sapiens* (human) | 284340 | NM_198477 | NP_940879 | MKVLISSLLLLLPLMLMSMVSSSLNPGVARGHRDRGQASRRWLQEG<br>GQECECKDWFLRAPRRKFMTVSGLPKKQCPCDHFKGNVKKTRHQRH<br>HRKPNKHSRACQQFLKQCQLRSFALPL |
| | Dr-CXCL17 | *Danio rerio* (zebrafish) | 100151367 | NM_001144821 | NP_001138293 | MKTMNFQILVLAFAVMIVTNIQCEARPQEGKSDKSAEVKGHAMPRK<br>CNCQVRGTALDRNCVCEMPHKSRPTLNPEQKNMCLKKKIKTFRKCL<br>QFMGANKKIAKGASLPI |
| | Dr-CXCL17-like | *Danio rerio* (zebrafish) | 100536854 | NM_001386806<br>XM_073906074 | NP_001373735<br>XP_073762175 | MTKPICLVFALLILTIILCNNSVCSQRRSMKQSAVCGCKLYPDKGL<br>KCTKRPNPKSRDEYYEILKCICRDTQIFSKSSRKEYLKRCNKFYPS<br>LPL |
| | Lc-CXCL17a | *Latimeria chalumnae* (coelacanth) | 106704074 | XM_014490303 | XP_014345789 | MKVSDVLVLLCAFTLSSCLQNSGSSDQEEGRVPEEASTVGAVKPQA<br>GSCSCGDPVGSLQNRLVSLKAGPSKHCDCRRSLKEAVNIKRPQAWA<br>WKSKPSRKRCKRKGDNGKIQIWGCRRKTIKPIIIPIPA |
| | Lc-CXCL17b | *Latimeria chalumnae* (coelacanth) | 106704074 | XM_014490304 | XP_014345790 | MKVSDVLVLLCAFTLSSCLQNSGSSDQEEGRVPEEASTVGAVKPQA<br>GSCSCGDPVGSLQNRLVSLKAGPSKHCDCRRSLKEVNIKRPQAWAW<br>KSKPSRKRCKRKGDNGKIQIWGCRRKTIKPIIIPIPA |
| GPR25 orthologs | Hs-GPR25 | *Homo sapiens* (human) | 2848 | NM_005298 | NP_005289 | MAPTEPWSPSPGSAPWDYSGLDGLEELELCPAGDLPYGYVYIPALY<br>LAAFAVGLLGNAFVVWLLAGRRGPRRLVDTFVLHLAAADLGFVLTL<br>PLWAAAAALGGRWPFGDGLCKLSSFALAGTRCAGALLLAGMSVDRY<br>LAVVKLLEARPLRTPRCALASCCGVWAVALLAGLPSLVYRGLQPLP<br>GGQDSQCGEEPSHAFQGLSLLLLLLTFVLPLVVTLFCYCRISRRLR<br>RPPHVGRARRNSLRIIFAIESTFVGSWLPFSALRAVFHLARLGALP<br>LPCPLLLALRWGLTIATCLAFVNSCANPLIYLLLDRSFRARALDGA<br>CGRTGRLARRISSASSLSRDDSSVFRCRAQAANTASASW |
| | Dr-GPR25 | *Danio rerio* (zebrafish) | 795188 | XM_073916757 | XP_073772858 | MASSTEMAHSGITMSLTSEYDYDYPINSTDENPIYTLPDAELLPMS<br>NIYIPVLYIIMFLTGSLGNLFVIVVIGKRRKKSGRLVDTFVLNLAL<br>ADLVFVLTLPMWAISTRYDEWPFGEVLCKISSFIIAVNRFSNIFFL<br>TCMSVDRYLAVVRLMDSRFLRSSNCAQITCGIVWVVSFFLGSPSLA<br>YRHLINNSVCSEDSKSSFVQGMNLLTILLTFLLPVLILGLCYGSIL<br>VNLRRHCHNPANTRTDARRRHSVKIVFAIISAFLISWLPFNCFKAI<br>HVALLIINGDLNEDTYVVIHRGLMLSCCLAFLNSCVNPAIYFFLDQ<br>HFRRRASMLCLSCLSQNDQAHQSYITSNSYSNGTSETCSGNTSTRG<br>RLFSLTQKA |
| | Lc-GPR25 | *Latimeria chalumnae* (coelacanth) | 102365624 | XM_005988473 | XP_005988535 | MPTESLQTASHDPSDFDYNADYSNFSTEDCDGDLPYAKIYIPIFYF<br>VIFFTGLFGNVFVIAAMTLKQTTKRLVDIFVINLAVADLVFVFTLP<br>LWSVSAAFDDQWLFGGVLCKLSSYVIAVNRYSSIFFMTGMSVDRYM<br>AVVKLLDSKFIRTRRCILITCTIIWIISLVMGIPSLVYRDLSTQDS<br>EHTYCIEDQDSIIFKGISLASLFLAFVLPVIIILFCYCSISARLYS<br>HFHHANRYDQKRKKTLKIIFTIITAFVCSWLPFNTFKTLYLLFSFQ<br>GKMPPCRVGILRQGLTITACFAFLSSCVNPIIYTFLDNHFRKRAHR<br>LLVKALGRYTERRNSFGESWASETSSTFVSRIRANSVKELQNMNKT<br>QQNTIPT |

**Table S2.** Summary of possible interactions of the coelacanth CXCL17a and CXCL17b with the coelacanth GPR25 predicted by AlphaFold3 algorithm. The binding structures of mature Lc-CXCL17a or Lc-CXCL17b with Lc-GPR25 were predicted via the online AlphaFold3 server (https://alphafoldserver.com).

| Ligand | Residues in ligand | Interacting residues in Lc-GPR25 |
|---|---|---|
| Lc-CXCL17a | A130 (carboxyl moiety) | R122 (TMD3) |
| | P129 | I118 (TMD3); ε-amine of K267 (TMD6) forms hydrogen bonds with carboxyl oxygen of P129 |
| | I128 | F263 (TMD6); F299 (TMD7) |
| | P127 | W94 (TMD2) |
| | I126 | L291 (TMD7) |
| Lc-CXCL17b | A129 (carboxyl moiety) | R122 (TMD3) |
| | P128 | I118 (TMD3); ε-amine of K267 (TMD6) forms hydrogen bonds with carboxyl oxygen of P129 |
| | I127 | F263 (TMD6); F299 (TMD7) |
| | P126 | W94 (TMD2) |
| | I125 | L291 (TMD7) |

**6×His-Lc-CXCL17a-3×Arg**

```
1   CTT TAA GAA GGA GAT ATA ATG CAT CAT CAC CAC CAT CAC CTG CAG AAC TCA GGC AGT TCT GAT CAG GAA GAA GGC
    GAA ATT CTT CCT CTA TAT TAC GTA GTA GTG GTG GTA GTG GAC GTC TTG AGT CCG TCA AGA CTA GTC CTT CTT CCG
                             M   H   H   H   H   H   H   L   Q   N   S   G   S   S   D   Q   E   E   G

76  CGT GTG CCA GAA GAG GCC TCT ACT GTT GGT GCG GTG AAA CCG CAA GCT GGC AGC TGC TCT TGT GGT GAC CCT GTT
    GCA CAC GGT CTT CTC CGG AGA TGA CAA CCA CGC CAC TTT GGC GTT CGA CCG TCG ACG AGA ACA CCA CTG GGA CAA
     R   V   P   E   E   A   S   T   V   G   A   V   K   P   Q   A   G   S   C   S   C   G   D   P   V

151 GGT TCA CTG CAA AAT CGT CTT GTC TCC CTG AAG GCG GGG CCG TCT AAA CAC TGT GAT TGT CGC CGT AGC TTG AAA
    CCA AGT GAC GTT TTA GCA GAA CAG AGG GAC TTC CGC CCC GGC AGA TTT GTG ACA CTA ACA GCG GCA TCG AAC TTT
     G   S   L   Q   N   R   L   V   S   L   K   A   G   P   S   K   H   C   D   C   R   R   S   L   K

226 GAA GCA GTA AAC ATC AAA CGT CCT CAG GCC TGG GCT TGG AAA TCC AAA CCG AGT CGC AAG CGC TGC AAG CGT AAA
    CTT CGT CAT TTG TAG TTT GCA GGA GTC CGG ACC CGA ACC TTT AGG TTT GGC TCA GCG TTC GCG ACG TTC GCA TTT
     E   A   V   N   I   K   R   P   Q   A   W   A   W   K   S   K   P   S   R   K   R   C   K   R   K

301 GGC GAC AAC GGG AAA ATT CAG ATT TGG GGT TGC CGT CGC AAA ACC ATC AAG CCA ATC ATC ATT CCG ATC CCT GCG
    CCG CTG TTG CCC TTT TAA GTC TAA ACC CCA ACG GCA GCG TTT TGG TAG TTC GGT TAG TAG TAA GGC TAG GGA CGC
     G   D   N   G   K   I   Q   I   W   G   C   R   R   K   T   I   K   P   I   I   I   P   I   P   A

376 CGT CGC CGT TAA GCG GCC GCA CTC GAG CAC CAC
    GCA GCG GCA ATT CGC CGG CGT GAG CTC GTG GTG
     R   R   R   *
```

**6×His-Lc-CXCL17b-3×Arg**

```
1   CTT TAA GAA GGA GAT ATA ATG CAT CAT CAC CAC CAT CAC CTG CAG AAC TCA GGC AGT TCT GAT CAG GAA GAA GGC
    GAA ATT CTT CCT CTA TAT TAC GTA GTA GTG GTG GTA GTG GAC GTC TTG AGT CCG TCA AGA CTA GTC CTT CTT CCG
                             M   H   H   H   H   H   H   L   Q   N   S   G   S   S   D   Q   E   E   G

76  CGT GTG CCA GAA GAG GCC TCT ACT GTT GGT GCG GTG AAA CCG CAA GCT GGC AGC TGC TCT TGT GGT GAC CCT GTT
    GCA CAC GGT CTT CTC CGG AGA TGA CAA CCA CGC CAC TTT GGC GTT CGA CCG TCG ACG AGA ACA CCA CTG GGA CAA
     R   V   P   E   E   A   S   T   V   G   A   V   K   P   Q   A   G   S   C   S   C   G   D   P   V

151 GGT TCA CTG CAA AAT CGT CTT GTC TCC CTG AAG GCG GGG CCG TCT AAA CAC TGT GAT TGT CGC CGT AGC TTG AAA
    CCA AGT GAC GTT TTA GCA GAA CAG AGG GAC TTC CGC CCC GGC AGA TTT GTG ACA CTA ACA GCG GCA TCG AAC TTT
     G   S   L   Q   N   R   L   V   S   L   K   A   G   P   S   K   H   C   D   C   R   R   S   L   K

226 GAA GTA AAC ATC AAA CGT CCT CAG GCC TGG GCT TGG AAA TCC AAA CCG AGT CGC AAG CGC TGC AAG CGT AAA GGC
    CTT CAT TTG TAG TTT GCA GGA GTC CGG ACC CGA ACC TTT AGG TTT GGC TCA GCG TTC GCG ACG TTC GCA TTT CCG
     E   V   N   I   K   R   P   Q   A   W   A   W   K   S   K   P   S   R   K   R   C   K   R   K   G

301 GAC AAC GGG AAA ATT CAG ATT TGG GGT TGC CGT CGC AAA ACC ATC AAG CCA ATC ATC ATT CCG ATC CCT GCG CGT
    CTG TTG CCC TTT TAA GTC TAA ACC CCA ACG GCA GCG TTT TGG TAG TTC GGT TAG TAG TAA GGC TAG GGA CGC GCA
     D   N   G   K   I   Q   I   W   G   C   R   R   K   T   I   K   P   I   I   I   P   I   P   A   R

376 CGC CGT TAA GCG GCC GCA CTC GAG CAC CAC
    GCG GCA ATT CGC CGG CGT GAG CTC GTG GTG
     R   R   *
```

**6×His-SmBiT-Lc-CXCL17a-3×Arg**

```
1   CTT TAA GAA GGA GAT ATA ATG CAT CAT CAC CAC CAT CAC GGT GTG ACC GGC TAC CGT CTG TTT GAA GAA ATT CTG
    GAA ATT CTT CCT CTA TAT TAC GTA GTA GTG GTG GTA GTG CCA CAC TGG CCG ATG GCA GAC AAA CTT CTT TAA GAC
                             M   H   H   H   H   H   H   G   V   T   G   Y   R   L   F   E   E   I   L

76  GGC GGC CTG CAG AAC TCA GGC AGT TCT GAT CAG GAA GAA GGC CGT GTG CCA GAA GAG GCC TCT ACT GTT GGT GCG
    CCG CCG GAC GTC TTG AGT CCG TCA AGA CTA GTC CTT CTT CCG GCA CAC GGT CTT CTC CGG AGA TGA CAA CCA CGC
     G   G   L   Q   N   S   G   S   S   D   Q   E   E   G   R   V   P   E   E   A   S   T   V   G   A

151 GTG AAA CCG CAA GCT GGC AGC TGC TCT TGT GGT GAC CCT GTT GGT TCA CTG CAA AAT CGT CTT GTC TCC CTG AAG
    CAC TTT GGC GTT CGA CCG TCG ACG AGA ACA CCA CTG GGA CAA CCA AGT GAC GTT TTA GCA GAA CAG AGG GAC TTC
     V   K   P   Q   A   G   S   C   S   C   G   D   P   V   G   S   L   Q   N   R   L   V   S   L   K

226 GCG GGG CCG TCT AAA CAC TGT GAT TGT CGC CGT AGC TTG AAA GAA GCA GTA AAC ATC AAA CGT CCT CAG GCC TGG
    CGC CCC GGC AGA TTT GTG ACA CTA ACA GCG GCA TCG AAC TTT CTT CGT CAT TTG TAG TTT GCA GGA GTC CGG ACC
     A   G   P   S   K   H   C   D   C   R   R   S   L   K   E   A   V   N   I   K   R   P   Q   A   W

301 GCT TGG AAA TCC AAA CCG AGT CGC AAG CGC TGC AAG CGT AAA GGC GAC AAC GGG AAA ATT CAG ATT TGG GGT TGC
    CGA ACC TTT AGG TTT GGC TCA GCG TTC GCG ACG TTC GCA TTT CCG CTG TTG CCC TTT TAA GTC TAA ACC CCA ACG
     A   W   K   S   K   P   S   R   K   R   C   K   R   K   G   D   N   G   K   I   Q   I   W   G   C

376 CGT CGC AAA ACC ATC AAG CCA ATC ATC ATT CCG ATC CCT GCG CGT CGC CGT TAA GCG GCC GCA CTC GAG CAC CAC
    GCA GCG TTT TGG TAG TTC GGT TAG TAG TAA GGC TAG GGA CGC GCA GCG GCA ATT CGC CGG CGT GAG CTC GTG GTG
     R   R   K   T   I   K   P   I   I   I   P   I   P   A   R   R   R   *
```

**Fig. S1.** The nucleotide sequence and amino acid sequence of the coelacanth CXCL17 precursors overexpressed in *E. coli*. The amino acid sequence of the mature Lc-CXCL17a or Lc-CXCL17b is shown in red, that of SmBiT in blue.

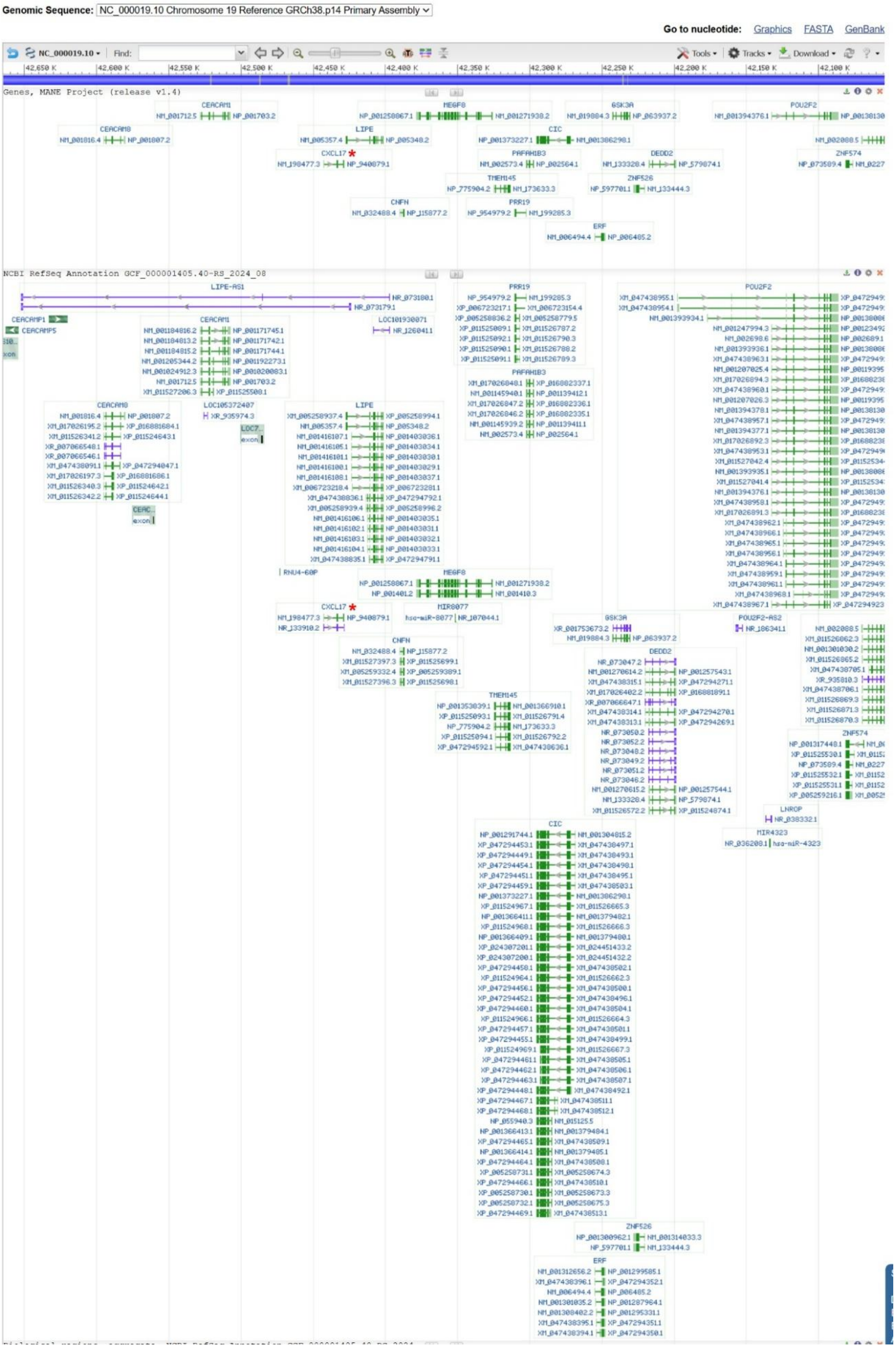


**Fig. S2.** Position of the human *CXCL17* gene in human genome. The human *CXCL17* gene is indicated by a red asterisk. The information was downloaded from the NCBI gene database (https://www.ncbi.nlm.nih.gov/gene/284340).

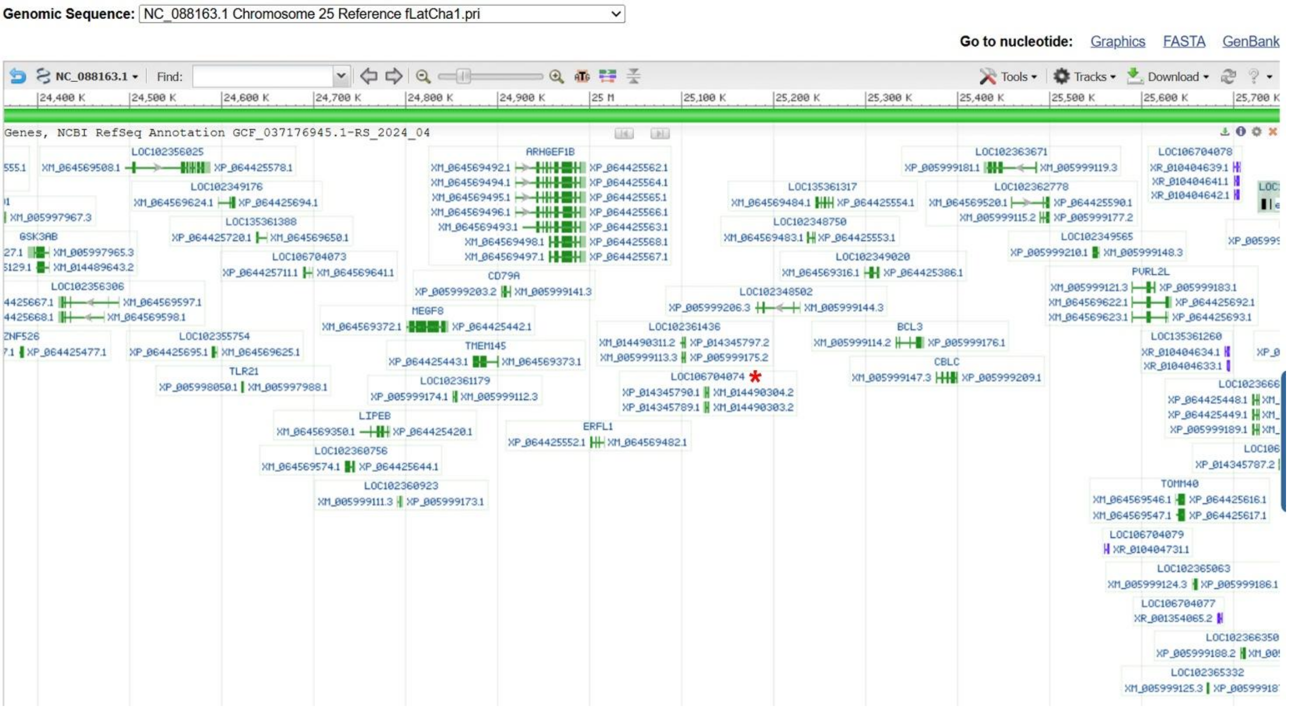


**Fig. S3.** Position of the coelacanth *CXCL17* gene in the genome of *Latimeria chalumnae*. The coelacanth *CXCL17* gene is indicated by a red asterisk. The information was downloaded from the NCBI gene database (https://www.ncbi.nlm.nih.gov/gene/?term=TMEM145%2C+coelacanth).

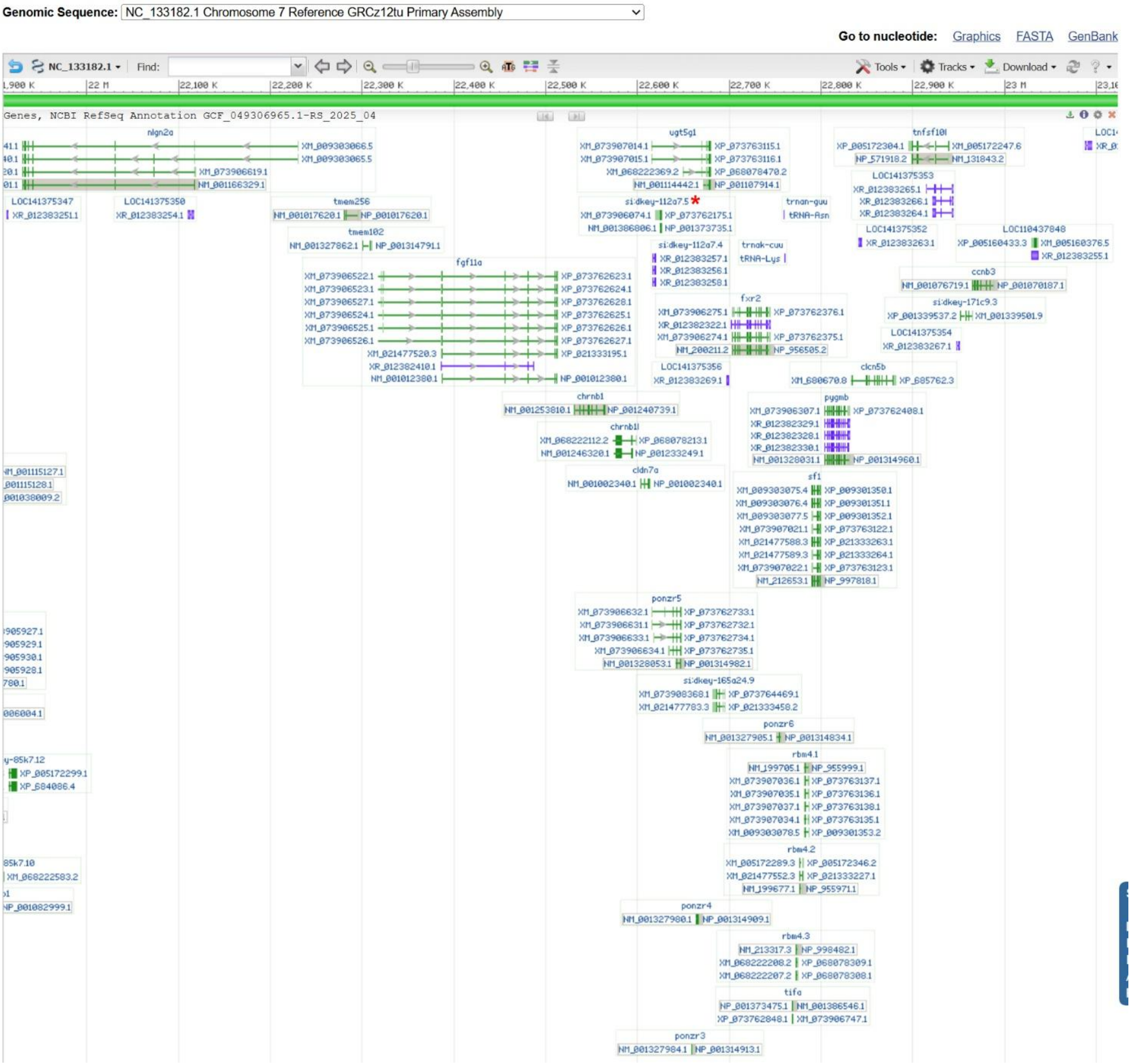


**Fig. S4.** Position of the zebrafish *cxcl17-like* gene in the genome of *Danio rerio*. The zebrafish *cxcl17-like* gene is indicated by a red asterisk. The information was downloaded from the NCBI gene database (https://www.ncbi.nlm.nih.gov/gene/?term=100536854).

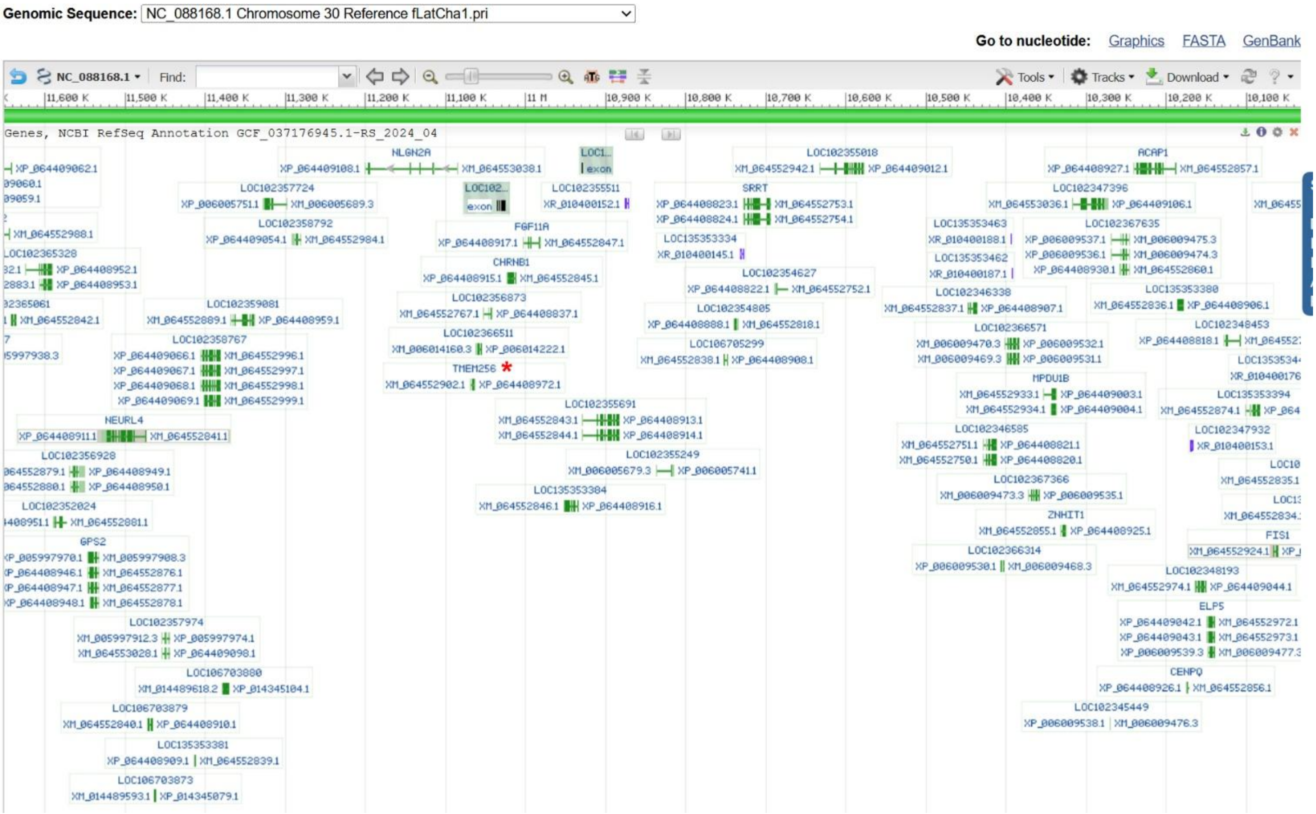


**Fig. S5.** Genes near to *TMEM256* in the genome of *Latimeria chalumnae*. The coelacanth *TMEM256* gene is indicated by a red asterisk. The information was downloaded from the NCBI gene database (https://www.ncbi.nlm.nih.gov/gene/?term=tmem256%2C+coelacanth).
